# Understanding spatiotemporal effects of food supplementation on host-parasite interactions using community-based science

**DOI:** 10.1101/2022.06.02.494595

**Authors:** Sarah A. Knutie, Rachel Bahouth, Matthew A. Bertone, Caroline Webb, Mahima Mehta, Mia Nahom, Rachael M. Barta, Sharan Ghai, Ashley C. Love, Sydney Horan, Alexandria Soldo, Elizabeth Cochrane, Jenna Bartholomew, Emily Cowan, Heather Bjerke, Susan L. Balenger, Michael W. Butler, Allison Cornell, Ashley C. Kennedy, Virginie Rolland, Elizabeth M. Schultz, Mark Stanback, Conor C. Taff, Gregory F. Albery

**Affiliations:** Department of Ecology and Evolutionary Biology, University of Connecticut, Storrs, CT, USA; Institute for Systems Genomics, University of Connecticut, Storrs, CT, 06269, USA; Department of Entomology and Plant Pathology, North Carolina State University, Raleigh, NC, USA 27695; Department of Biology, University of Mississippi, University, MS, USA; Department of Biology, Lafayette College, Easton, PA, 18042, USA; Department of Biology, Penn State Altoona, Altoona, Pennsylvania, USA; Mosquito Control Section, Delaware Division of Fish and Wildlife, Newark, DE, 19702, USA; Department of Biology, Arkansas State University, AR, 72467, USA; Wittenberg University, Department of Biology, Springfield, OH, 45504, USA; Department of Biology, Davidson College, Davidson, NC, 28035, USA; Lab of Ornithology and Department of Ecology & Evolutionary Biology, Cornell University, Ithaca, NY, 14850, USA; Department of Biology, Georgetown University, Washington, DC, 20007, USA

**Keywords:** Citizen science, disease ecology, ectoparasites, food supplementation, host-parasite interactions

## Abstract

1. Supplemental feeding can increase the overall health of animals but also can have variable effects on how animals defend themselves against parasites. However, the spatiotemporal effects of food supplementation on host-parasite interactions remain poorly understood, likely because large-scale, coordinated efforts are difficult.
2. Here, we introduce the Nest Parasite Community Science Project, which is a community-based science project that coordinates studies with bird nest box “stewards” from the public and scientific community. This project was established to understand broad ecological patterns between hosts and their parasites.
3. The goal of this study was to determine the effect of food supplementation on eastern bluebirds (*Sialia sialis*) and their nest parasite community across the geographic range of the bluebirds from 2018–2021. We received 646 nests from 68 stewards in 26 states in the eastern United States. Nest box stewards reported whether or not they fed their bluebirds mealworms or suet, then followed the nesting success of the birds (number of eggs laid and hatched, percent hatched, number and percent fledged). We then identified and quantified parasites in the nests.
4. We found that food supplementation increased fledgling numbers and proportional fledging success. The main nest parasite taxa were parasitic blow flies (*Protocalliphora sialia*), but a few nests contained fleas (*Ceratophyllus idius, C. gallinae, Orchopeas leucopus*) and mites (*Dermanyssus* spp. and *Ornithonyssus* spp.). Blow flies were primarily found at northern latitudes, where food supplementation affected blow fly abundance. However, the direction of this effect varied substantially in direction and magnitude across years. More stewards fed bluebirds at southern latitudes than at northern latitudes, which contradicted the findings of other community-based science projects.
5. Overall, food supplementation of birds was associated with increased host fitness but did not appear to play a consistent role in defense against these parasites across all years. Our study demonstrates the importance of coordinated studies across years and locations to understand the effects of environmental heterogeneity, including human-based food supplementation, on host-parasite dynamics. Studies during a single year or considering only a single population might not provide the necessary data to develop management strategies for species.

## Introduction

Environmental factors, such as food availability, can influence host-parasite interactions (Becker et al., 2015, 2018; Sánchez et al., 2018). Host defense strategies against parasites, such as resistance, are often condition-dependent and affected by food availability. Resistance reduces the damage that parasites cause by reducing parasite fitness (Read et al., 2008). Resistance mechanisms, such as the immune response, can be condition-dependent because mounting these responses can be energetically costly (Howick & Lazzaro, 2014; Knutie, 2020; Lochmiller & Deerenberg, 2000; Sheldon & Verhulst, 1996). Therefore, only hosts with enough food resources are able to invest in a robust immune response. Extra nutrients directly increase immune cell production, which may account for the positive relationship between food availability and immunity (Strandin et al., 2018). For example, supplemented protein can increase the concentration of cellular and humoral immune cells (Coop & Kyriazakis, 2001; Datta et al., 1998). Consequently, food availability is expected to influence parasite abundance, but evidence for this phenomenon in the wild is mixed.

Humans can change resource availability for animals by intentionally providing food using wild bird feeders or unintentionally by leaving food waste in the environment (Murray et al., 2016). In fact, humans provide many wild bird species with a large proportion of their food (Cox & Gaston, 2018; Jones, 2011). In the United States alone, approximately 50 million households provide over half a million tons of supplemental food to attract wild birds to their property (Cox & Gaston, 2016; Robb et al., 2008). Supplemental feeding of birds can have several benefits to birds and humans. Feeding wild birds can improve the mental health of humans and strengthen their connection with nature (Cox & Gaston, 2016, 2018; Jones, 2011; Shaw et al., 2017). Birds that are supplemented with food are often in better condition, which, in turn, can increase their reproductive success (Bailey & Bonter, 2022; Tollington et al., 2019) and enhance some measures of immunity (Cornelius Ruhs et al., 2019; Lochmiller et al., 1993; Sánchez et al., 2018; Strandin et al., 2018; Wilcoxen et al., 2015).

Recent experimental work with a wild bird species demonstrated that food supplementation increases resistance to parasitism, but this study relied on only one year of data in one population (Knutie, 2020). Due to environmental heterogeneity, studies are needed across years and populations to understand the broad impact of food supplementation on host-parasite interactions. Such studies are difficult to accomplish without coordinated efforts, such as community-based science projects (e.g., eBird, NestWatch; Phillips & Dickinson, 2009; Sullivan et al., 2009). These projects have provided a wealth of data to understand the impact of factors, including food supplementation, on bird fitness (Bailey & Bonter, 2022). However, these studies have limitations because they cannot provide insight into interspecific interactions, such as host-parasite relationships. Thus, the Nest Parasite Community Science Project (hereafter “Project”), was established. This community-based science project works with the public and scientific community (hereafter “stewards”) to monitor bird nest boxes. Generally, the Project explores the effect of environmental conditions on spatiotemporal patterns of box-nesting birds, such as eastern bluebirds (*Sialia sialis*), and their nest parasite community.

The eastern bluebird (hereafter “bluebird”) is a North American bird species that is supplemented with food by humans. In the 1970s, populations of bluebirds declined, in part because of a loss of suitable foraging and nesting habitat (Gowaty & Plissner, 2020). In response, humans built and established artificial nest boxes and some began supplementing the birds’ natural diet of insects, spiders, and small fruits (Pinkowski, 1977) with dried and live mealworms (larvae of *Tenebrio molitor*). Since the 1970s, the bluebird population size rebounded within approximately a decade (Sauer & Droege, 1990) and humans continue to maintain nest boxes and provide bluebirds with supplemental food. These continued efforts are likely important, as bluebirds face challenges, such as parasitic, nest-inhabiting flies, throughout much of their range (Grab et al., 2019). Past studies have found that blow fly (Calliphoridae: *Protocalliphora*) abundances are highly variable, and these flies either have a negative effect or no effect on fledging success of bluebirds (reviewed in Grab et al., 2019). A recent study in Minnesota found that bluebirds supplemented with mealworms had higher resistance (via an antibody response) to blow flies than unsupplemented birds (Knutie, 2020). However, even within populations, blow fly abundances and effects on survival were highly variable across years (Grab et al., 2019).

The goal of this study was to determine the effect of food supplementation on host-parasite interactions across years and geographic locations. Nest box stewards either fed their bluebirds mealworms and/or suet or not, then followed the nesting success of the birds (number of eggs laid and hatched, proportion of eggs hatched, number of fledglings, proportion of nestlings fledged) across the geographic range of the bluebird from 2018–2021. Once the nests were empty, they were sent to the University of Connecticut and we identified and quantified nest parasite taxa. First, we used this information to determine spatial and temporal effects on nesting success and nest parasite presence and abundance. Second, we determined the effect of food supplementation on fledging success and parasite prevalence. Lastly, because stewards voluntarily did or did not feed their bluebirds mealworms, we determined whether there was a spatial effect to bird feeding by stewards.

## Methods

### Field methods

Nest box stewards were recruited from 2017-2021 through social media platforms (e.g., Twitter and Facebook groups). From 2018–2021, we received a total of 674 nests from 68 stewards across 26 states in the eastern United States (Table S1). These stewards noted whether they provided no food, mealworms (live or dried), or suet to bluebirds on their property. Suet contains animal fat and other items such as corn meal, peanuts, fruits, and/ or dried insects. Forty stewards from 21 states provided mealworms in at least one of the years and 38 stewards from 19 states did not; seven stewards from seven states provided mealworms in some years but not the other years. The exact number of mealworms provided to the bluebirds varied by the steward. Stewards noted that they added 50–200 mealworms per day to the feeders, which were 0–27 m (mean = 12 m) from the nest boxes.

Stewards were instructed to remove any old material from boxes in March-April each year. Stewards then monitored their nest boxes based on when bluebirds were expected to arrive on the breeding grounds (e.g., March for more southern latitudes and May for more northern latitudes). Once a nest box had nesting material, stewards confirmed that the nest box was occupied by bluebirds. The number of eggs laid in the box were counted visually once the clutches were complete. Once the eggs hatched, the stewards visually counted the number of nestlings. The stewards monitored the survival of nestlings until the nest was empty or dead nestlings were present. After the nest was empty, stewards removed the nest from the box and placed it individually in a gallon-sized, plastic ziplock bag. They placed a labeled piece of paper in the bag with the following information: collection date, full name and steward identification number (ID), city, county, state, zip code, bird species, whether mealworms were fed, number of eggs laid, number of nestlings that hatched, and number of nestlings that died. We also calculated the proportion of nestlings that survived until fledging (i.e. fledging success). If infertile eggs or dead nestlings were found, stewards were instructed to remove these items with gloves before shipping the nests. Once the bags were labeled, stewards placed the bags in a cool, dry area. After the breeding season was complete, nests were compiled and shipped in a cardboard box or paper envelope to the University of Connecticut.

### Parasite identification, quantification, and measurements

After nests were received, they were logged into a database and assigned to a nest dissector. Immediately prior to dissection, the ziplock bag was placed in a -80 °C freezer for approximately 10 min (but sometimes up to an hour) to immobilize any live invertebrates. Once the nest was removed from the freezer, pieces of the nest material were removed from the bag and dissected over a white piece of paper, which took between 30 min to 2 h, depending on the number of invertebrates in the nest. All invertebrates were collected from the nest material and placed in 2 mL tubes with 90% ethanol. Specimens were then stored in a -80 °C freezer until they were identified. We were unable to dissect 28 nests because they were too wet to dissect and thus reliably find the smaller parasites.

Specimens were identified into broad taxonomic groups and then sent to MAB for identification. Blow fly pupae were identified under a dissecting scope after being removed from alcohol and dried; no further preparation was performed. Flea and mite specimens were slide-mounted in Hoyer’s mounting medium or by clearing first in 10% KOH, washing, and mounting in PVA (lactic acid, phenol, and polyvinyl alcohol); slides were left to cure on a slide warmer. Nests also contained commensal book lice (Liposcelididae), but since they were not parasitic we excluded them from the study.

Nests contained parasitic, commensal, and predatory mites and therefore we separated these groups before identifying the parasitic genera. Identifications were confirmed to major groups: commensal dust mites (Pyroglyphidae), predatory mites of other mites (Cheyletidae), and parasitic mites (Mesostigmata). Mesostigmata specimens were slide-mounted and identifications of the genera *Ornythonyssus* and *Dermanyssus* were made using a compound microscope at various magnifications (200–1000X) and using published diagnostic keys (Di Palma et al., 2012; Knee & Proctor, 2006; Murillo & Mullens, 2017). Non-parasitic (commensal and predatory) mites were excluded from the study. Flea identifications were made using a compound microscope at various magnifications (200–400X) and using published diagnostic keys (Holland, 1951, 1985; Lewis, 2000). Blow flies were identified using available morphological keys for pupae (Whitworth, 2003; Whitworth, 2003). We also measured the width of empty pupal cases for up to 10 individuals per nest as a proxy for fly size, which is related to lifetime fitness in Diptera (Moon, 1980; Schmidt & Blume, 1973). We measured 1004 pupal cases from 126 nests of 13 stewards. We could not measure pupal case length because flies emerge from the top of the case, thus removing part of it.

### Statistical analyses

We used generalized linear mixed models (GLMMs) to examine spatiotemporal drivers of the five fitness components and abundances of each parasite taxa. All analyses were conducted in R (version 4.1.1; R Core Team, 2023), using the integrated nested Laplace approximation (INLA; Lindgren & Rue, 2015). All models were checked by simulating from the posterior and verifying the even distribution of residuals and verifying that the models’ simulated data recapitulated the distribution of the input data.

#### Host fitness models

We fitted models that examined each of our five fitness metrics as response variables. We used a Gaussian distribution for the number of eggs laid and hatched and number of nestlings that fledged, and binomial distribution for the proportion of eggs that hatched and the proportion of nestlings that successfully fledged (i.e. fledging success). For the former, the number of eggs laid represented the number of trials and the number of eggs hatched represented the number of successes and for the latter, the number of nestlings represented the number of trials and the number of fledglings represented the number of successes. Explanatory variables included: year (categorical with four levels: 2018, 2019, 2020, 2021); and day-of-year that the nests were collected (1–365, representing number of days since January 1st). We included steward ID as a random effect to account for among-site variation in fitness. To ask whether supplementation improved fitness (i.e. for the number and proportion of nestlings fledged) we also fitted supplementation as a fixed effect. We did not include this effect for eggs laid and hatched and proportion hatched because supplementation was not always commenced before eggs were laid, meaning that we would not be able to reliably infer effects of supplementation on these components of fitness.

#### Parasite models

Parasite models used blow fly infection as a binary response variable; for the other parasites, prevalence was too low to fit a reliable model, both for fleas (2.8%) and for mites (3.7%). Fixed effects included supplementation, year, day-of-year, and number of eggs hatched, with steward ID as a random effect. When exploring the data, substantial among-year variation in the effects of supplementation on parasite prevalence was apparent; as such, we included supplementation as an interaction with year to examine the differences in the effect of supplementation on blow fly prevalence across years. This model showed stronger support (i.e., improved model fit when compared using DIC) than including separate main effects of supplementation. We also added some elaborations to this model: first, because there were strong spatial patterns in blow fly prevalence, we also repeated the model including only the latitudes above which these parasites had been found (above 39.7 °N), which is corroborated by past studies (Sabrosky et al., 1989). Second, we repeated the analysis with the highly overdispersed counts of blow fly abundances using a negative binomial distribution to investigate whether abundance showed the same trends as prevalence.

#### Blow fly size models

We also determined whether variation in blow fly size (pupal width [mm]) could be explained by any of the fixed effects. Fixed effects included year, year-by-supplementation interaction, number of eggs hatched, and total parasite abundance with nest and steward as random effects.

#### Spatial autocorrelation effects

For all fitness and parasite models, we fitted a stochastic partial differentiation equation (SPDE) effect to control for and quantify spatial autocorrelation in the response variable (Lindgren et al., 2011; Lindgren & Rue, 2015). The SPDE effect uses samples’ bivariate coordinates to model spatial dependence, examining whether samples from closer locations are more similar and then generating a two-dimensional spatial field that can be examined for spatial patterns. This approach has proved successful for investigating spatial patterns of parasite prevalence and intensity (Albery et al., 2019, 2022). We fitted an SPDE effect based on samples’ latitude and longitude and examined whether it improved model fit by assessing whether it reduced the deviance information criterion (DIC) of the model, using deltaDIC=2 as a cutoff. INLA also allows fitting of separate spatial fields for different time periods, and therefore we also included between-year variation in the spatial field and assessed whether it improved model fit using the same cutoff.

#### Steward models

Finally, we examined whether the probability of a steward providing food to their birds varied spatially. We fitted a model with the binary response variable of food supplementation (yes or no), and with fixed effects including year, the total number of nests submitted from the property, latitude, and longitude (all continuous). For all models, because there were two types of supplementation (mealworms and suet), we attempted to use each method on its own as an explanatory variable to investigate whether they had different implications for hosts and parasites. However, there were no notable differences among these specifications, and therefore we reported the fullest models – i.e., those combining mealworm and suet supplementation – alone.

## Results

### Fitness models

Bluebirds laid between 2–6 eggs that hatched between 0–6 nestlings (*n* = 354 nests). Our fitness models revealed strong effects of food supplementation on the number of fledglings (effect size 0.317, confidence intervals (0.138, 0.495); P<0.001), and proportion fledging success (effect size 0.071, confidence intervals (0.018, 0.126); *P* = 0.01; Figure 1A, S1C), but on no other host fitness variables (Figure S1C). We detected strong spatial heterogeneity in the distribution of two out of five fitness metrics. For number of fledglings, there was a patchy distribution with alternating hot- and coldspots (Figure 1C; ΔDIC=-6.58), and contrastingly, the east coast had a higher proportional fledging success than inland birds (Figure 1D; ΔDIC=-15.17). All other metrics did not show a significant improvement when the spatial effect was included (Figure S1; ΔDIC>-2). In addition, our models revealed a number of day-of-year effects (Figure S1; all *P* < 0.001). Nests sampled later in the year had fewer eggs (-0.335, CI (-0.414, -0.256)) and fewer hatchlings (-0.223, CI (-0.305, -0.142)), but had a greater proportional fledgling success (0.036, CI (0.014, 0.057)).

**Fig. 1.**
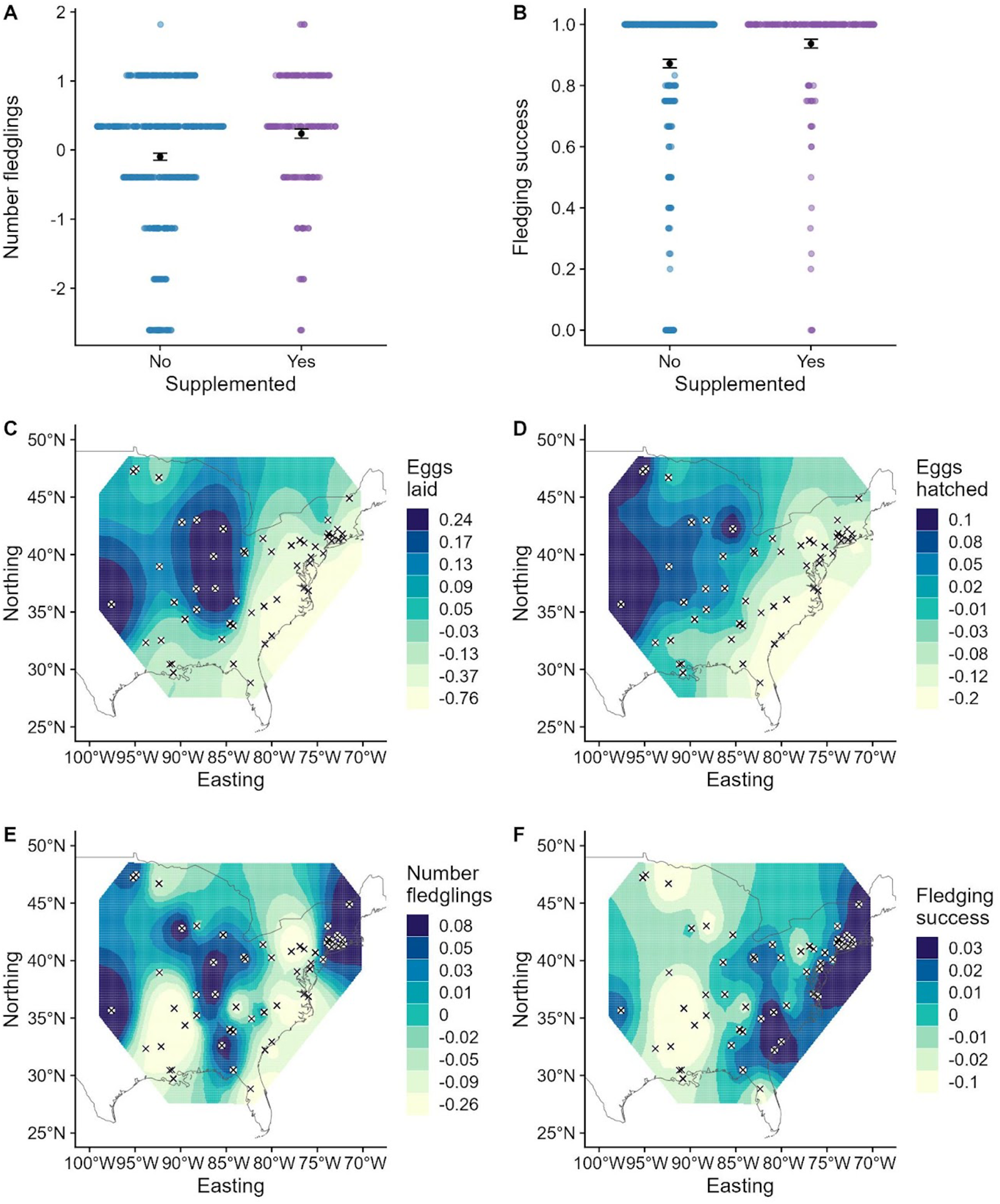
Eastern bluebird fitness variables were affected by A-B) food supplementation and C-F) geographic location. In Panels A-B, each coloured point represents a nest; the black dot represents the mean, and the error bars denote the standard error around this estimate. The number of fledglings (A) was transformed to have a mean of 0 and a standard deviation of 1. Panels C-F display the geographic distribution of the spatial random effect for numbers of eggs laid, eggs hatched, number of fledglings, and proportional fledging success. Points with crosses represent sampling locations. Darker colors represent greater numbers of eggs laid (C) and hatched (D) and number of fledglings (E) in units of standard deviations, and proportion fledging success (F) on the logit scale.

### Parasite models

Out of 646 nests that were dissected for parasites, 171 nests (26.5%) from 20 stewards across 11 states contained 1–139 blow flies, which were all identified as *Protocalliphora sialia*. Eighteen nests (2.8%) from seven stewards across six states contained 1– 179 fleas. Most flea taxa were identified as *Ceratophyllus idius*, but one nest contained *C. gallinae* and one nest contained *Orchopeas leucopus*. Twenty-four nests (3.7%) from 10 stewards across nine states contained between one to over 4000 parasitic mites from the genera *Dermanyssus* and *Ornithonyssus*.

We found strong support for spatial effects in blow fly prevalence (ΔDIC = -5.80); inspecting the projected spatial effects, prevalence decreased from North to South (Figures 2B). Notably, blow flies were only found as far south as 39.77°N (Figure 2B). We uncovered strong support for among-year variable supplementation effects on blow fly abundance (ΔDIC = -5.56; Figure 2A, S2). The effect of supplementation on blow fly abundance was positive in 2018, negative in 2020, and not significantly different in 2019 and 2021 (Figure 2A). Additionally, there was a substantial positive effect of day-of-year on blow fly abundance (0.681, CI (0.3, 1.069), *P* < 0.001; Figure S2).

**Fig. 2.**
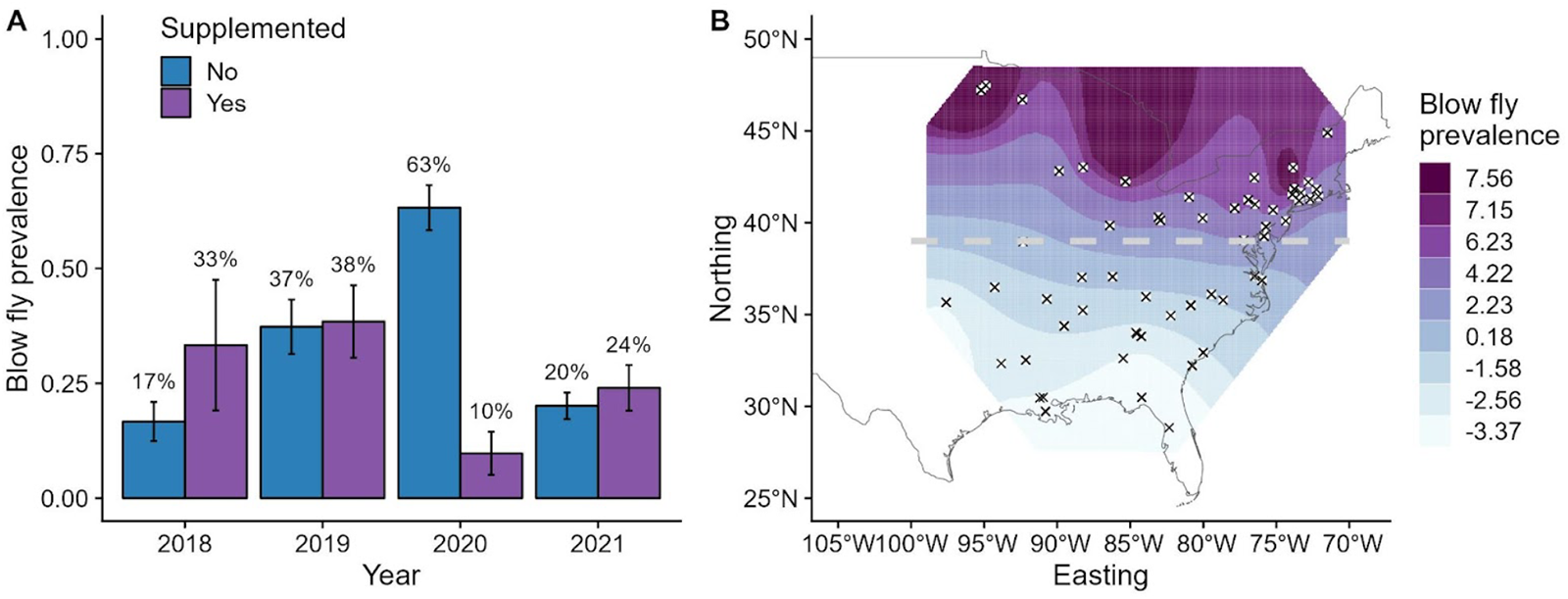
Parasite prevalence (i.e., the proportion of nests with blow flies) was affected by food supplementation in different years (A) and geographic location (B). Panel A presents the relationship between supplementation (yes or no; x-axis) and blow fly prevalence (y-axis). The proportion of nests with blow flies for supplemented birds compared to unsupplemented birds was higher in 2018, similar in 2019, lower in 2020, and similar in 2021. The percentage of nests with parasites is displayed above the bars. Panel B displays the geographic distribution of the spatial random effect for blow fly prevalence (B). Points with crosses represent sampling locations. Darker colors represent greater parasite prevalence on the logit scale. The grey dashed line represents the latitude below which no blow flies were found.

Blow fly size did not have strong spatial effects (ΔDIC > 2, Figure S3). Year and supplementation did not significantly affect blow fly size (Figure S3), but number of eggs hatched and blow fly size correlated positively (0.157, CI (0.034, 0.28), *P* = 0.013).

### Steward models

Finally, our steward models uncovered a negative correlation between stewards’ latitude and their probability of feeding their birds mealworms (-0.891, CI (-1.401, – 0.381); *P* < 0.001). That is, stewards at more southern latitudes were substantially more likely to feed their birds than those at more northern latitudes (Figure 3, S4).

**Fig. 3.**
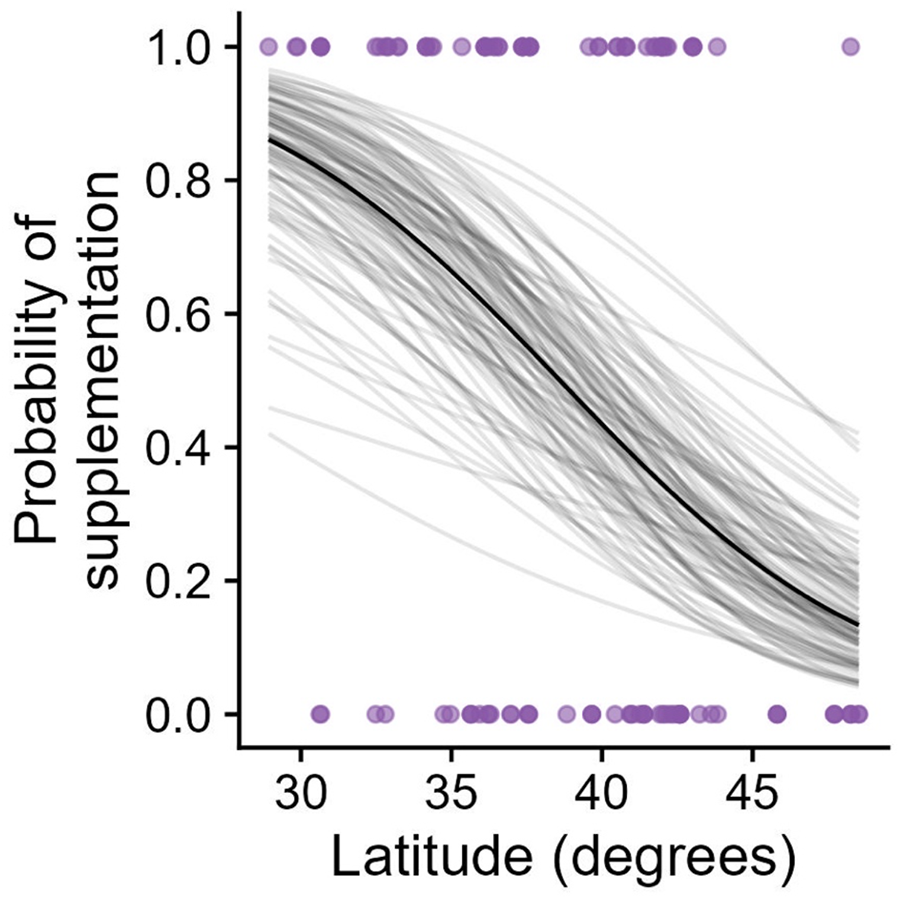
Stewards in higher latitudes were substantially less likely to feed birds on their property. Points represent an individual steward in a given year and the grey lines represent 100 random draws from the posterior distribution of our model estimates for the effect of latitude, which displays uncertainty in the estimate. The black line represents the mean of this distribution, thereby showing the mean effect estimate.

## Discussion

Our study introduces the Nest Parasite Community Science Project from which we assessed the effect of food supplementation on eastern bluebirds and their nest parasite community across years and the geographic range of bluebirds. We found that the number of eggs laid and hatched were higher in the midwestern United States than on the east coast, but this effect was not observed for hatching success or fledging success. Food supplementation increased fledging success, which has also been shown for bluebirds in a recent study (Bailey & Bonter, 2022). The main nest parasite taxa were parasitic blow flies, fleas, and mites. Blow flies were found only in the northern latitudes, as observed in previous studies (reviewed in Sabrosky et al., 1989). Fleas and mites were more rare in bluebird nests and fleas were only found in the northern latitudes. Within the range of the blow flies, food supplementation affected blow fly abundance but this effect varied across years. Finally, more stewards fed bluebirds at southern latitudes than at northern latitudes. Our results suggest that host-parasite dynamics can vary spatiotemporally, including in response to food supplementation of the host.

On average, bluebirds in the Midwest laid and hatched more eggs, with numbers decreasing moving towards the East Coast. The average number of eggs laid and hatched within a population can be constrained by food availability and energetic constraints, and by the number of offspring that the parents can feed (Food Limitation Hypothesis; Lack, 1947). Therefore, food might be more abundant in the midwest than on the East Coast, allowing for females to lay and hatch more eggs. The proportion of nestlings that hatched did not differ spatially, implying that there was not a similar geographic effect on egg fertility or viability.

Fleas (*Ceratophyllus* spp.) and blow flies (*Protocalliphora sialia*) were found only in northern latitudes. Decades of past work have also shown that the geographical range of *Protocalliphora* spp. is restricted to northern latitudes (Sabrosky et al., 1989). These patterns might be because ectotherm survival is dependent on environmental temperature (Munch & Salinas, 2009). Parasites were only found in areas where they experience temperate winter conditions, such as freezing temperatures. In turn, these insects have evolved strategies to deal with winter conditions. For example, shorter days and lower temperatures can signal insects to enter into a state of developmental arrest, called diapause, to survive (Vinogradova & Reznik, 2013). During diapause, reduced metabolic rate can slow down the process of aging and increase survival (Denlinger, 2008; Keil et al., 2015). For example, higher temperatures can reduce diapause survival in butterflies (*Lycaena tityrus*) and the rice stem borer (*Chilo suppressalis*) (Klockmann & Fischer, 2019; Xiao et al., 2017). Thus, parasites might be unable to survive the higher winter temperatures in the south for long enough to take advantage of the next host breeding season. Higher summer temperatures at southern latitudes might also reduce the fitness of fleas and blow flies. Mean summer temperature can negatively predict nest parasite load (Musgrave et al., 2019). Temperature-dependent production of pigments like AGE, which is a marker of cellular senescence, can accumulate more quickly at higher temperatures in the related brown blow fly (*Calliphora stygia*; Kelly et al., 2013). Experimental studies are needed to determine how seasonal temperatures affect the survival and evolution of fleas and blow flies, which would then causally explain the patterns observed in our study.

Interestingly, we found that few bluebird nests contained mites, and those that contained mites included the genera *Dermanyssus* and *Ornithonyssus.* These two genera have been found in the nests of many species of passerines and on the bodies of domestic poultry, such as chickens (Proctor et al., 2000). In passerine nests, mites either overwinter in old nests and then infest new nests the following year, or they transmit into the nests on nesting material or adult birds (Proctor et al., 2000). Tree swallows, which are another common box-nesting bird, are often highly infested by parasitic nest mites (Knutie, unpublished data; Winkler, 1993). This species often incorporates chicken feathers into their nests, which not only provides the opportunity for mites to invade the nests, but also results in higher mite abundance (Winkler, 1993). Bluebird or mite behavior might alternatively explain the lack of mites in bluebird nests. For example, bluebirds might choose boxes that are not apparently infested by mites, as found in other species with their roost sites (Christe et al., 1994). Mites might prefer other hosts compared to bluebirds if bluebirds have suboptimal characteristics (e.g., small brood sizes; Burtt Jr. et al., 1991) or effective defenses against mites (e.g., preening, immune response). The differing infestation rates among box-nesting host species deserves further attention and would provide more insight into multi-host-parasite dynamics (Albert et al., 2023; Grab et al., 2019).

Food supplementation had contrasting effects on blow fly abundance in different years. In 2018 supplemented nests had more parasites than non-supplemented nests, in 2019 and 2021 there was no difference between treatments, and in 2020 supplemented bird nests had fewer parasites. One possible driver for these patterns is variation in food availability from natural sources. Food supplementation can improve host immune responses (Tschirren et al., 2007), and experimental work has demonstrated that supplemented bluebirds invest more in resistance mechanisms, which reduces parasite abundance (Knutie, 2020). However, this effect was primarily observed only early in the season when resource availability was low. Thus, immune investment by bluebirds might have varied across years due to changing food resource availability. For example, aerial insect abundance and activity can increase with temperature (Winkler et al., 2013), so temperature differences across years could have resulted in differing food (insect) availability. Blow flies themselves might be responding to changes in annual temperatures, with survival and fecundity changing in response to dynamic winter or summer temperatures. Although mean blow fly abundance in supplemented nests remained constant, abundances did vary across years in unsupplemented nests. Overall, our results suggest that annual variation in environmental conditions likely affects both host defenses and blow fly fitness. Furthermore, these results suggest that environmental effects can vary among years and provides further evidence that researchers must consider that a single year of data might not provide the whole story. Characterizing the effect of other environmental factors on host-parasite interactions is beyond the scope of this study, but can be explored further with this community-based science project.

Close to 50 million Americans feed birds in their backyards or around their homes (Cox & Gaston, 2016; Robb et al., 2008). We found that stewards fed bluebirds in the southern United States more than in the north (Fig. 3), which contradicts other studies. For example, several studies have found that people from northern latitudes feed birds more than those in the south (Horn & Johansen, 2013). Systematic reporting biases could potentially explain our results. For example, southern bird-feeding enthusiasts or hobbyists are more likely to volunteer in community science projects than their northern counterparts (Lepczyk et al., 2012). Furthermore, our contradictory spatial patterns could be specific to bluebirds, given that none of the aforementioned studies focused exclusively on bluebirds. Spatial biases are common in citizen science projects (Geldmann et al., 2016) and can be explained by human population density. For example, higher population density, as found at northern latitudes (United States Census Bureau, 2022), correlates with higher sampling efforts. Future studies could focus on explaining why bluebirds are fed by more stewards in the southern US than in the northern US, which could have conservation implications for the species.

Community-based science projects can provide a wealth of data that can help researchers explore broad spatial and temporal ecological patterns that might not otherwise be possible. The main result of our study demonstrates that food supplementation can have varying effects on host-parasite interactions across years and thus, cautions the interpretation of results from only one year of data. Additionally, we found that fleas and blow flies are restricted to the northern geographic range of bluebirds, which begs the question of whether southern bluebirds, which are not infested with fleas or blow flies, have evolved the same immune defenses against ectoparasites as northern bluebirds (Knutie, 2020). Given the amount of training and permits involved in handling animals for research, a coordinated study on the evolution of host immune differences across geographic areas would only be possible with the academic scientists involved in the Nest Parasite Community Science Project.

## Supporting information

Supplemental Results

## Acknowledgments

We thank the following nest box stewards for their participation in the study: Nancy McDonald, Gigi Gerben, Denise Hartranft, Elsabe Allen, Pattie Barnes, Annette Evans, Tammy C. Brown, Doug Finck, Paula Chambers, Sally Grames, Max Wilson, John Hurlbert, Amy Wray, George Mitchell, Steve Dingeldein, Honey Winger, Heather Harris, Bobbie Moller, Angela Tedesco, Aubrie Giroux, Diane Hutchins, Wanda Carlock, Harrison Ramey, Jan Raia, Deanna Gott, Sandi and Steve Knutie, Laurie Hanson, Patrice Murphy, Dawn Denson, Rory Larson, Deb Houck, Janet Allison, Cindi and Bill Kobak, Kat Smith, Susan Gilnack, Patty Roberts, Alan Biggs, Matt Daley, Rochelle Bowman, Kim Klaproth, Debby Martin, Paula Robertson, Hilda Young, Jimmie Espich, Kathy Kerns, Tamara “Toma” Yermakova-Nikolayenko, Joe and Sue Brinson, Martha Lackman, Cathy Chiovaro, Karen Barnes, Krista Tschaenn, Sarabeth Klueh-Mundy, Joe Bear, Ava Fontenot, Lori Conley, Erin Kuprewicz, Carlos Garcia Robledo, Brooke Mikles, Mary McGregor, Suzi Litaker, Becky Boyd, Maya Stine, Grace Emin, Julianna Carpenetti, Emma Stierhoff, Jessica Gutierrez, Lauren Albert, Vel Johnston. We appreciate graduate and undergraduate students who assisted with nest monitoring. Thanks to the Knutie lab for feedback on the manuscript (Alyssa McGurer, Lorraine Perez, Heather Skeen). The work was supported by a National Science Foundation CAREER grant (IOS-2143899) and University of Connecticut (UConn) start-up funds to SAK and a UConn Summer Undergraduate Research Fellowship to RB. GFA was supported by a College for Life Sciences Fellowship from the Wissenschaftskolleg zu Berlin, and by NSF grant number DEB-2211287. All applicable international, national, and/or institutional guidelines for the care and use of animals were followed. All bird handling and work was conducted according to approved UConn IACUC protocols (No. A18-005) with the appropriate state permits or waivers and U.S. Fisheries and Wildlife Service Scientific Collecting Permit #MB11631D. All raw data will be available on FigShare upon acceptance. Authors declare no conflict of interest.

## Statement on Inclusion

Our study brings together authors throughout the United States, where the study took place. All authors were engaged early on with the research and study design to ensure that the diverse sets of perspectives they represent were considered from the onset and undergraduate researchers participated in the writing of the discussion.

## Author contributions

SAK and RB designed the project, SAK, RB, MAB, CW, MM, MN, RMB, SG, ACL, SH, AS, EC, JB, EC, HB, SB, MWB, AC, AK, VR, EMS, MS, and CCT collected data, GFA conducted analyses, MAB, SEH, and AES identified parasites, SAK, MAB, RB, CW, MM, MN, RMB, SG and GFA wrote the manuscript and all authors revised the manuscript.

