## Supplemental Results for "Understanding spatiotemporal effects of food supplementation on host-parasite interactions using community-based science"

**Supplemental material. Knutie et al.**

**Appendix 1.** Understanding spatiotemporal effects of food supplementation on host-parasite interactions using community-based science.

Table S1. The number of nests from landlords based on state and year.

| State | 2018 | 2019 | 2020 | 2021 | **Total** |
|---|---|---|---|---|---|
| AL | 0 | 0 | 1 | 4 | **5** |
| AR | 0 | 29 | 0 | 25 | **54** |
| CT | 2 | 3 | 3 | 9 | **17** |
| DE | 10 | 0 | 0 | 0 | **10** |
| FL | 2 | 4 | 3 | 2 | **11** |
| GA | 5 | 7 | 1 | 3 | **16** |
| IN | 0 | 3 | 2 | 3 | **8** |
| KY | 0 | 3 | 5 | 3 | **11** |
| LA | 0 | 0 | 7 | 4 | **11** |
| MA | 0 | 0 | 0 | 1 | **1** |
| MD | 0 | 2 | 5 | 8 | **15** |
| MI | 0 | 9 | 11 | 12 | **32** |
| MN | 15 | 34 | 38 | 17 | **104** |
| MO | 0 | 0 | 1 | 0 | **1** |
| MS | 42 | 0 | 0 | 0 | **42** |
| NC | 0 | 6 | 0 | 110 | **116** |
| NH | 0 | 2 | 2 | 2 | **6** |
| NJ | 0 | 0 | 4 | 0 | **4** |
| NY | 1 | 8 | 34 | 15 | **58** |
| OH | 2 | 5 | 3 | 6 | **16** |
| OK | 2 | 3 | 3 | 3 | **11** |

| | | | | | |
|---|---|---|---|---|---|
| PA | 8 | 12 | 16 | 31 | **67** |
| SC | 0 | 0 | 8 | 15 | **23** |
| TN | 2 | 0 | 3 | 4 | **9** |
| VA | 0 | 1 | 4 | 8 | **13** |
| WI | 11 | 0 | 2 | 0 | **13** |
| **Total** | **102** | **131** | **156** | **285** | **674** |

Supplementary Figure 1: DIC values associated with model fits (top row) and fixed effect estimates (bottom row) for the fitness variable models. In the bottom row, dots represent the mean of the posterior distribution of the effect; the error bars represent the 95% credibility intervals of the effect.

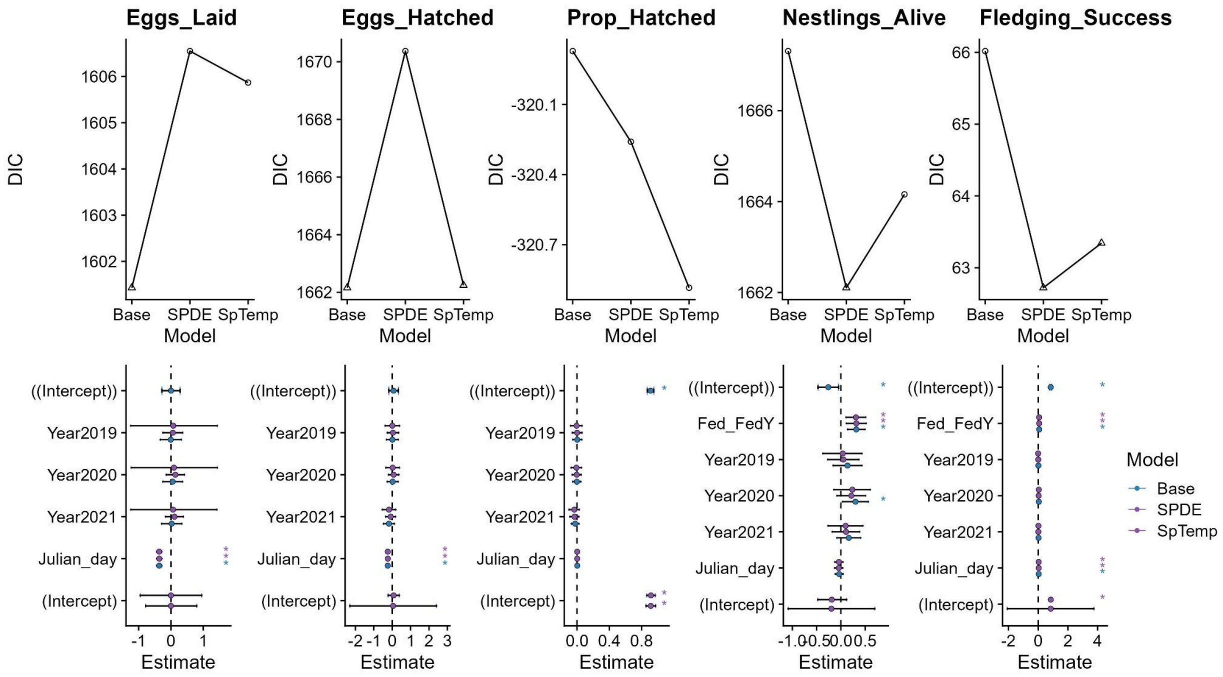

Supplementary Figure 2: DIC values associated with model fits (top row) and fixed effect estimates (bottom row) for the blow fly prevalence models. In the bottom row, dots represent the mean of the posterior distribution of the effect; the error bars represent the 95% credibility intervals of the effect.

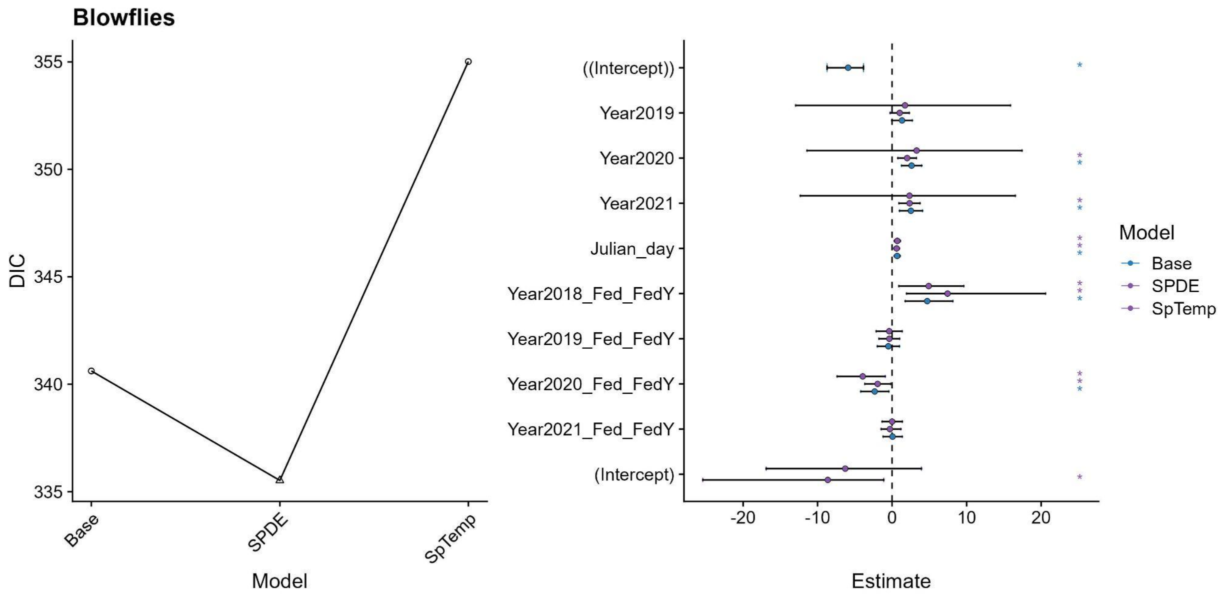

Supplementary Figure 3: DIC values associated with model fits (left panel) and fixed effect estimates (right panel) for the parasite size models. In the right panel, dots represent the mean of the posterior distribution of the effect; the error bars represent the 95% credibility intervals of the effect.

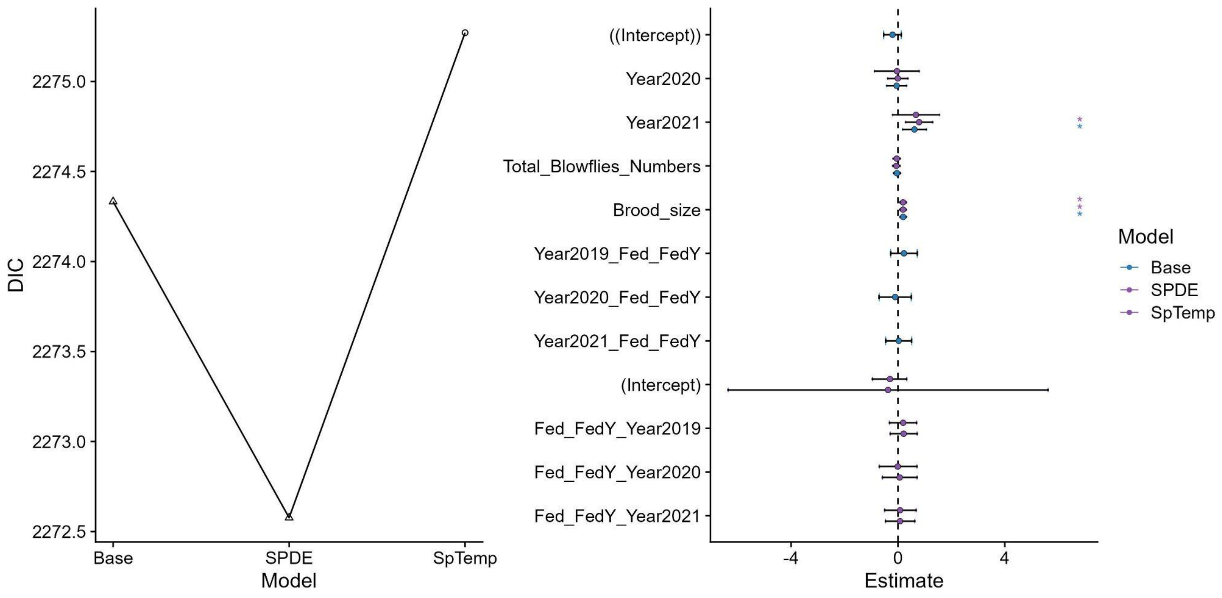

Supplementary Figure 4: DIC values associated with model fits (left panel) and fixed effect estimates (right panel) for the landlord supplementation models. In the right panel, dots represent the mean of the posterior distribution of the effect; the error bars represent the 95% credibility intervals of the effect.

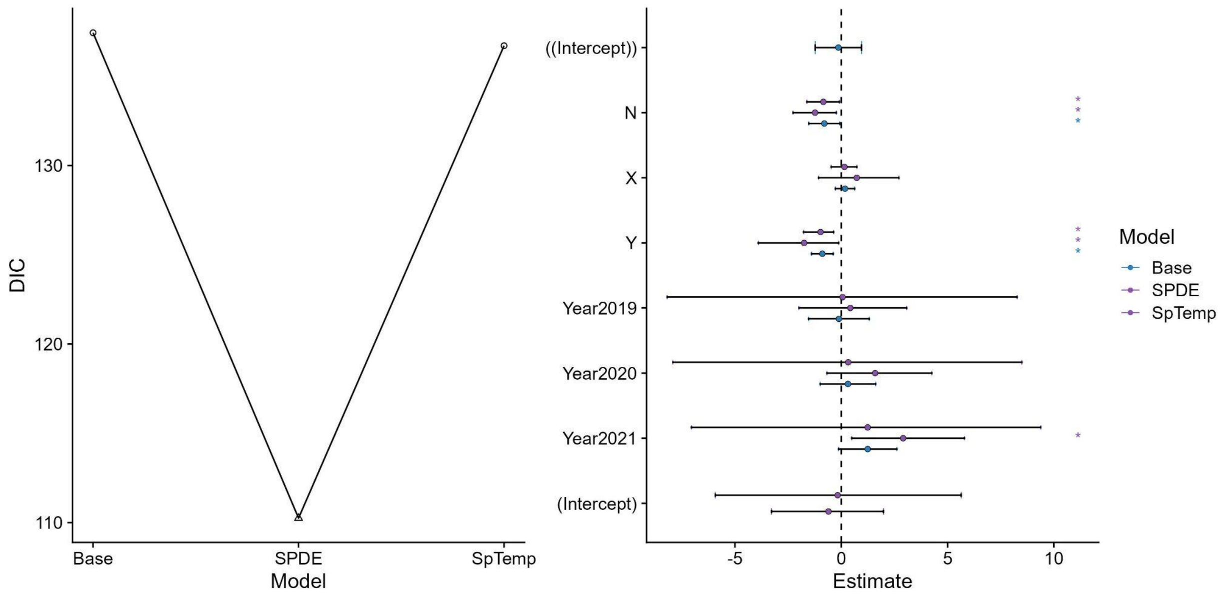
